# Perinatal folate levels do not influence tumor latency or multiplicity in a model of NF1 associated plexiform-like neurofibromas

**DOI:** 10.1101/2023.01.13.523962

**Authors:** Andrew R. Marley, Kyle B. Williams, Justin Tibbitts, Christopher L. Moertel, Kimberly J. Johnson, Michael A. Linden, David A. Largaespada, Erin L. Marcotte

## Abstract

**Objective:** In epidemiological and experimental research, high folic acid intake has been demonstrated to accelerate tumor development among populations with genetic and/or molecular susceptibility to cancer. Neurofibromatosis type 1 (NF1) is a common autosomal dominant disorder predisposing affected individuals to tumorigenesis, including benign plexiform neurofibromas; however, understanding of factors associated with tumor risk in NF1 patients is limited. Therefore, we investigated whether pregestational folic acid intake modified plexiform-like peripheral nerve sheath tumor risk in a transgenic NF1 murine model.

**Results:** We observed no significant differences in overall survival according to folate group. Relative to controls (180 days), median survival did not statistically differ in deficient (174 days, *P*=0.56) or supplemented (177 days, *P*=0.13) folate groups. Dietary folate intake was positively associated with RBC folate levels at weaning, (*P*=0.023, 0.0096, and 0.0006 for deficient vs. control, control vs. supplemented, and deficient vs. supplemented groups, respectively). Dorsal root ganglia (DRG), brachial plexi, and sciatic nerves were assessed according to folate group. Mice in the folate deficient group had significantly more enlarged DRG relative to controls (*P*=0.044), but no other groups statistically differed. No significant differences for brachial plexi or sciatic nerve enlargement were observed according to folate status.

## Introduction

Neurofibromatosis type 1 (NF1), a disorder caused by inherited or *de novo NF1* gene mutations, is characterized by the development of benign neurofibromas, occurring in 25-30% of all NF1 patients [1–4]. As these neurofibromas can be congenital [2], perigestational exposures may influence their development. Among increased risk for several other malignancies [5–8], malignant peripheral nerve sheath tumors (MPNSTs) are the most common malignant NF1-associated tumors (10-13% lifetime risk of occurrence) [9, 10]. As malignant transformation from benign plexiform neurofibromas to MPNST has been proposed [11], those with benign plexiform neurofibromas have an increased risk of developing MPNSTs relative to NF1 patients without such benign tumors [12, 13]. Transformation to MPNST is of particular concern, as these have no targeted therapies available for treatment and patients have poor 5-year survival [14]. Clinical manifestations of NF1 vary [15], and little is known regarding factors that influence tumor risk among NF1-affected individuals.

We hypothesized that maternal folic acid intake during pregnancy may influence offspring tumor risk. In the general population, increased adult folate intake has been found to decrease risk of several malignancies [16–20], and epidemiological evidence has suggested an inverse association between peri-gestational maternal folic acid intake and risk for disease outcomes in offspring [21–25]. In contrast, adults with ‘genetic susceptibility’, pre-neoplastic lesions, or who are at risk of cancer recurrence may have an increased risk for tumorigenesis when combined with high folate intake [26–29]. Experimental studies in rodents [30–33] have largely corroborated these findings. Collectively, these studies suggest that, in the presence of genetic and/or molecular susceptibility, folate supplementation may promote tumor growth. However, these mechanisms have yet to be experimentally explored in the context of MPNST risk among NF1 patients.

We sought to address this literature gap by determining whether perigestational folic acid exposure modifies risk of plexiform-like peripheral nerve sheath tumors in transgenic NF1 mouse models [34]. Our objective was to understand whether varying folate intake in a mouse model of NF1 associated plexiform-like neurofibromas during the perigestational period modulates the incidence and severity of tumor formation in the offspring, hypothesizing that reduced maternal folic acid intake during pregnancy decreases rate and severity of tumor formation in offspring. Our results may inform future human clinical trials regarding optimal maternal dietary folate intake in populations genetically susceptible to cancer syndromes.

## Methods

### Animals and Diets

The University of Minnesota Institutional Animal Care and Use Committee (IACUC) and the US Army Medical Research and Material Command (USAMRMC) Animal Care and Use Review Office (ACURO) approved all protocols for this study. Mice were housed in plastic cages layered with shavings in a temperature and humidity-controlled room. Mice had 12-hour day/night cycles with access to food and water *ad libitum*. Mice used for breeding included *Dhh*-Cre; *Nf1^flox/flox^* males and *Nf1^flox/flox^*; *pten^flox/flox^* females. One month prior to mating, 6–10-week-old female mice were assigned to one of three aminoacid defined diets, containing 0.3 mg/kg (deficient), 2.0 mg/kg (control), or 8.0 mg/kg (supplemented) folic acid and 1% succinyl sulfathiazole (SST) (Harlan Teklad, Madison, WI: catalogue numbers TD.07056, TD.07057, and TD.07058) to prevent folic acid synthesis by intestinal microflora [35]. There were 25, 15, and 5 dams assigned to the deficient, control, and supplemented folic acid groups, respectively. Due to fecundity concerns in folate-deficient mice, all dams were switched to standard diet for a month after weaning their first litter then returned to experimental diet before producing a second litter. Female mice were harem-mated at two-to-three per cage with one *Dhh*-Cre; *Nf1^flox/flox^* male mouse. Female mice were maintained on their respective diets until three weeks post-delivery, at which time pups were weaned and switched to standard chow for the remainder of the experiment. In conjunction with the University of Minnesota Research Animals Resources staff and veterinary technicians, mice were monitored for neurological indications of tumor development, such as ataxia and limb paralysis. Animals were euthanized by CO_2_ inhalation and cervical dislocation in accordance with institutional guidelines at 12 months (specified endpoint), tumor formation, or when moribund for any reason, whichever came first. Necropsies were conducted to assess the presence of enlarged per-herbal nerves and potential tumor burden. Abnormal sciatic nerves and brachial or sacral plexi were also removed. Enlarged dorsal root ganglia (DRG) were counted for the entire spinal cord. Figure 1 schematically summarizes our experimental design and procedures.

**Figure 1.**
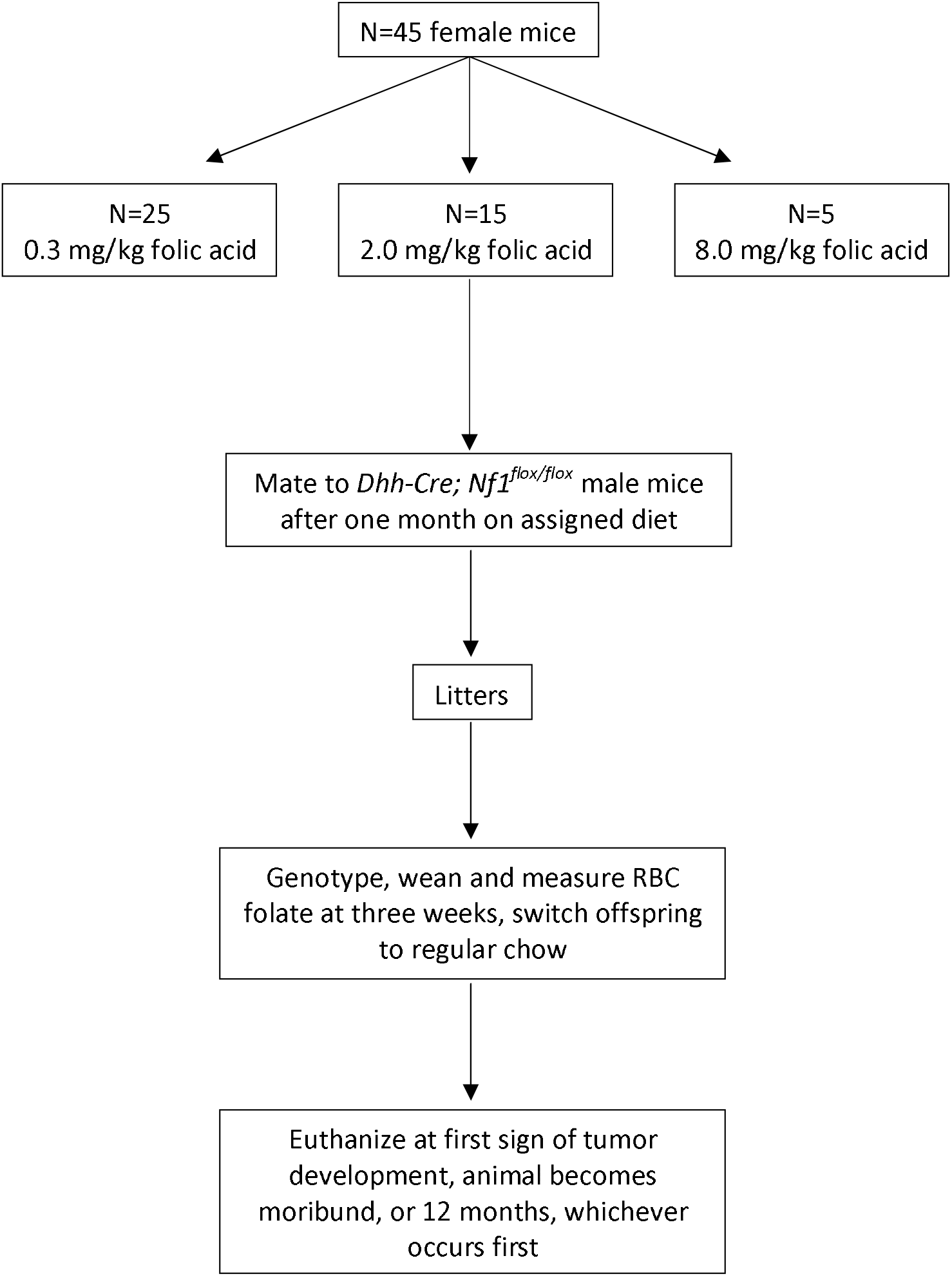
Experimental design for study of triple transgenic *Nf1* mice assigned to folic acid supplementation groups to observe tumor growth and survival in offspring.

### Genotyping and Red Blood Cell (RBC) Folate Measurement

Full genotyping and RBC folate measurement methods can be found in the supplement.

### Statistical Analysis

Based on our hypothesis that offspring of folate deficient mice will experience increased survival and offspring of supplemented mice will experience decreased survival relative to controls, we chose an unbalanced design (allocating more mice to folate deficient diets and fewer mice to supplemented diets) to maximize statistical efficiency. To examine the effect of folic acid on rate of tumor development, we utilized Kaplan-Meier survival tests to assess whether time to tumor development varied according to dietary group. Survival time was calculated as number of days from birth to tumor development or one year of age, whichever came first. Differences in RBC folate and DRG were assessed using Brown-Forsythe and Welch’s analysis of variance (ANOVA) and Dunnett’s T3 test for multiple comparisons. Differences in sciatic nerve enlargement were assessed using Kruskal-Wallis ANOVA and Dunn’s test for multiple comparisons. Statistical analyses were performed using GraphPad Prism (version 9.4.1) (GraphPad Software, San Diego, CA). Tests were two-sided and p-values of <0.05 were considered statistically significant.

## Results

A total of 45 female mice were assigned to deficient (n=25), control (n=15), and supplemented (n=5) folate diets. Dams assigned to a deficient diet produced 88 (48.4%) of the 182 total experimental offspring. Corresponding numbers for control and supplemented folate dams were 81 out of 159 (50.9%), and 21 out of 47 (44.7%), respectively. There was no statistically significant difference in percent experimental offspring according to folate group (*P*=0.73). Kaplan-Meier survival analyses indicated no significant differences in overall survival according to folate group. Relative to controls (180 days), median survival did not statistically differ in deficient (174 days, *P*=0.56) or supplemented (177 days, *P*=0.13) folate groups (Figure 2). Maternal dietary folate intake was significantly, positively associated with RBC folate levels at weaning, as significant differences were observed among all group comparisons (*P*=0.023, 0.0096, and 0.0006 for deficient vs. control, control vs. supplemented, and deficient vs. supplemented groups, respectively) (Figure 3). DRG, brachial plexi, and sciatic nerves of 62 mice from the deficient group, 68 mice from the control group, and 20 mice from the supplemented group were examined via necropsy to assess tumor burden (Supplemental Figure 1). Mice from dams with a deficient, control, and supplemented folate diet had averages (SDs) of 4.02 (2.10), 3.16 (1.83), and 3.60 (1.64) enlarged DRG, respectively, with mice in the deficient group having significantly more enlarged DRG relative to controls (*P* = 0.044). Comparisons among other groups did not statistically differ. Brachial plexus enlargement did not differ according to folate status. Among folate deficient, control, and supplemented dams, 87.3%, 85.5%, and 78.6%, respectively, had enlarged brachial plexi (*P*=>0.99, >0.99, and 0.83 for deficient vs. control, control vs. supplemented, and deficient vs. supplemented groups, respectively. We also observed no significant difference in sciatic nerve enlargement among supplemented (20.0%), control (14.7%), or deficient (17.7%) groups (*P*=>0.99 for all comparisons).

**Figure 2.**
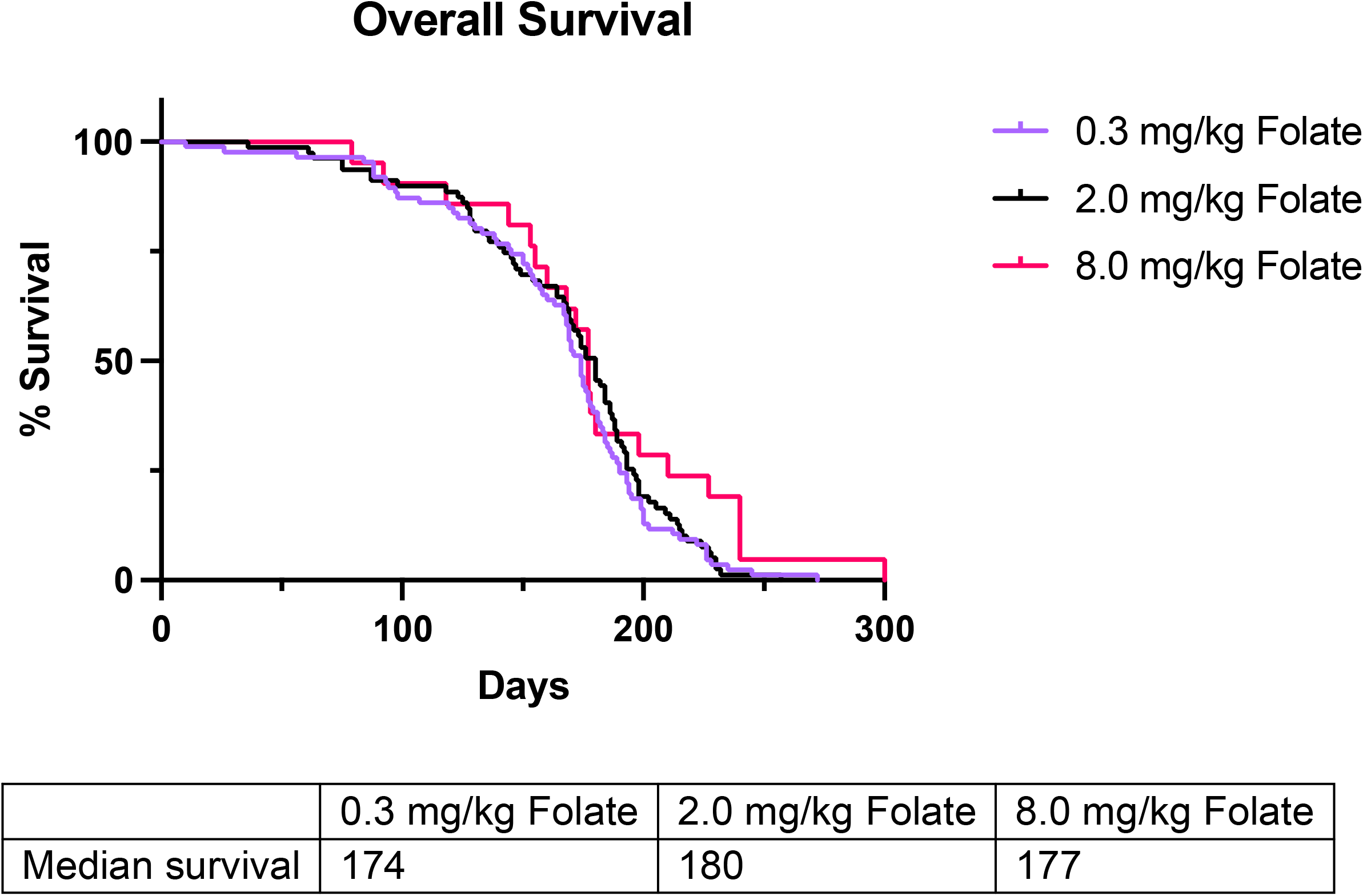
Overall survival in transgenic NF1 mouse model according to dietary folate consumption.

**Figure 3.**
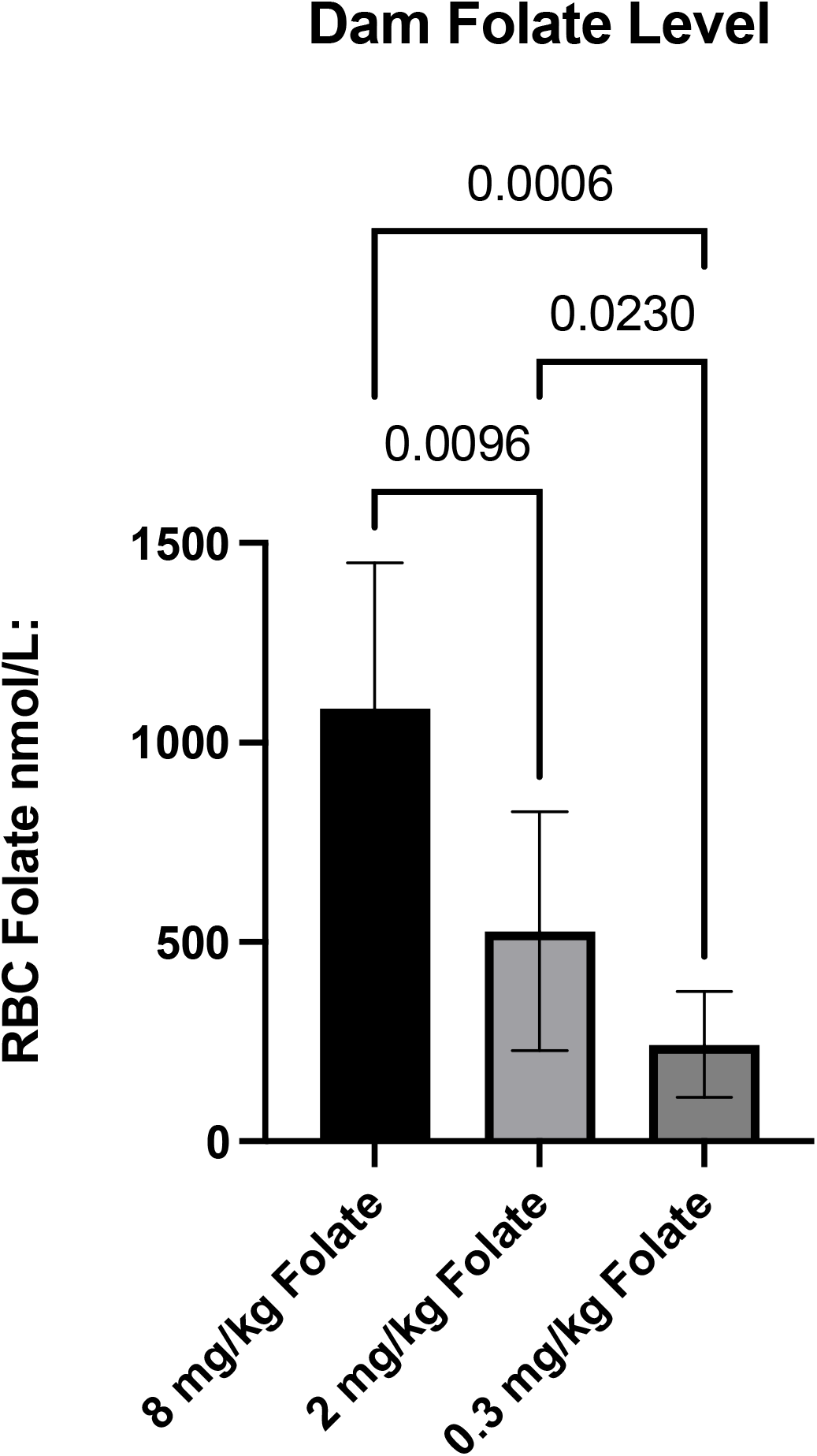
RBC folate level at weaning according to maternal dietary folate intake.

## Discussion

Although maternal folate intake is critical to healthy fetal growth and development [36, 37], folate may also fuel growth and proliferation of neoplastic cells [38–40], increasing risk of tumorigenesis in susceptible populations. Current results, however, do not corroborate this line of evidence, as folate supplementation did not affect overall survival in our susceptible population of NF1-affected mice.

In our previous analysis of a mouse model with nevoid basal cell carcinoma syndrome, offspring of dams assigned a low folate diet had reduced risk of medulloblastoma relative to control and a statistically significant increase in medulloblastoma survival [31]. Similarly, in a study of adenoma-susceptible mice, Lawrance *et al*. found folate deficiency prior to conception was associated with decreased adenomas, while high folate introduced after weaning was associated with increased adenomas [30]. Deghan Manshadi *et al*., in a study of mammary tumors initiated in female rats at puberty, demonstrated accelerated tumor progression and greater tumor weight/volume associated with folic acid supplementation 2.5x the control concentration [33]. Sharma *et al*. discovered that high folate progressed the development of hepatocellular carcinoma among rats given diethylnitrosamine carcinogen [41]. In a previous study also investigating nerve sheath tumors, mice carrying the human T-lymphotropic virus type 1 transactivator gene and provided high folate developed tumors significantly sooner than carrier mice assigned to low folate [32]. Among humans, a large randomized trial of folic acid supplementation among participants with a history of colorectal adenomas found supplementation was associated with increased risk of ≥3 adenomas [26], and a prospective analysis of women with *BRCA1/2* mutation found a positive association between plasma folate and breast cancer risk [42], Additional experimental and human studies also demonstrated risk associated with high folate both with and without known genetic or medical susceptibility [27–29, 43].

It is possible that our smaller sample may have contributed to the inconsistency with previous findings, as our current study assigned 25, 15, and 5 dams to deficient, control, and supplemented folate categories, respectively, versus the 42 dams assigned to respective categories in our previous study [31]. However, given our results, it is unlikely that a larger sample size would have made a meaningful impact, particularly considering the significant results observed in other studies with small samples [30, 32]. The inconsistency could also be due to differences in administered folate concentrations. In the current study, deficient, control, and supplemented diets contained 0.3, 2.0, and 8.0 mg/kg folic acid whereas the high folate group in Lawrance *et al*. received 20mg/kg folic acid [30], and Sharma *et al*. provided low and high folate groups with 0 mg/kg folate and 20 mg/kg folate, respectively [41]. The higher concentration of administered folate in supplemented groups and/or wider range of dietary consumption in other studies could help explain the inconsistency of our results. However, folate concentrations administered in the current study are identical to concentrations administered in our previous study [31], similar to concentrations administered by Deghan Manshadi *et al*. (2, 5, 8, and 10 mg/kg folic acid) [33], and greater than concentrations assigned by Bills et al. (0.11 micromoles/kg folic acid [~0.05 mg/kg] and 11.34 micromoles/kg folic acid [~5 mg/kg]) [32]. As all three of these studies demonstrated statistically significant findings, the inconsistency with our results is unlikely due to folate dosage. Also, because of the findings of Bills *et al*. [32] and the demonstration of folate-associated effects in similar tissues [44–46], the inconsistency of our results cannot likely be attributed to etiologic and/or biological differences between soft tissue sarcomas and other various tumor types. Therefore, current results may differ from previous analyses due to folate being administered perigestationally versus directly to offspring. Although previous findings [30, 31] demonstrated associations between maternal folate and adverse events in offspring [31], and it is established that the maternal intrauterine environment can impact offspring cancer risk [47–49], it is possible that maternal conditions are not as critical to plexiform-like neurofibroma formation relative to other tumors. Future studies on the effects dietary folate levels throughout life may reveal a role in peripheral nerve sheath tumor development.

### Conclusions

Our results failed to support the hypothesis that high perigestational folic acid intake in a model of NF1-affected mice increases the rate or severity plexiform-like tumor formation in offspring. Further study is needed to confirm these findings and assess the safety of high perigestational folic acid intake in the presence of cancer-susceptibility syndromes, particularly NF1.

## Limitations

Our study was limited by only being able to detect tumor initiation and progression through monitoring onset of neurological symptoms, motor function impairment, and paralysis in the mice. It is possible that imaging modalities, such as a longitudinal MRI study, could have shown differences in tumor initiation and/or growth rates that our study was unable to discern. Additionally, physiological and pathological differences in folate metabolism and neurofibroma progression between mice and humans limit our ability to extrapolate findings to human populations.

## Abbreviations

NF1: neurofibromatosis type 1
MPNST: malignant peripheral nerve sheath tumors
IACUC: The University of Minnesota Institutional Animal Care and Use Committee
USAMRMC: US Army Medical Research and Material Command
ACURO: Animal Care and Use Review Office
DRG: dorsal root ganglion
ANOVA: analysis of variance

## Declarations

### Ethics approval and consent to participate

Ethics approval for this study was provided by the University of Minnesota Institutional Animal Care and Use Committee and the US Army Medical Research and Material Command Animal Care and Use Review Office.

### Availability of data and material

Data used to support findings reported herein are available from the corresponding author upon request.

## Acknowledgements

Not applicable

## Consent for publication

Not applicable

## Authors’ contributions

EM, DL, and KJ conceptualized the study, ML and DL acquired the data, DL provided murine models and laboratory resources and oversaw study procedures, KW and JT performed laboratory tests and tended to the animals, KW performed data analyses, AM, CM, EM, KJ, ML, and JT interpreted the data, AM wrote and submitted the manuscript, all authors contributed to manuscript edits and revisions, and all authors read and approved the final manuscript.

