## Supplemental Materials for "Perinatal folate levels do not influence tumor latency or multiplicity in a model of NF1 associated plexiform-like neurofibromas"

**Table of Contents for Supplementary Materials**

**Genotyping methods.** Methods utilized in the genotyping of a transgenic NF1 murine model

**Red Blood Cell (RBC) Folate Measurement.** Description of methods used to measure RBC folate concentration.

**Supplementary Table 1.** (a) Number of dorsal root ganglia (DRG), (b) brachial plexi enlargement, and (c) sciatic nerve enlargement according to dietary folate intake.

*Genotyping Methods*

Genotyping was conducted at weaning by isolating DNA from tail clippings using standard proteinase K treatment, phenol-chloroform extraction, and ethanol precipitation. Genomic DNA was dissolved in sterile TE [10 mmol/L tris-HCL (7.5 pH), 1 mmol/L EDTA (8 pH)] and quantified using a Nanodrop spectrophotometer. PCR genotyping was conducted using 50ng of diluted genomic DNA as a template in a 25mL PCR reaction volume. PCR primers used for *Dhh-Cre* were forward 5’-CTGGCCTGGTCTGGACACAGTGCCC-3’ and reverse 5’-CAGGGTCCGCTCGGGCATAC-3’ (amplicon 385 bp); *Nf1* floxed allele were wild type (WT) forward 5’-CTTCAGACTGATTGTTGTAACTGA-3’, WT reverse 5’-ACCTCTCTAGCCTCAGGAATGA-3’, and floxed reverse 5’-TGATTCCCACTTTGTGGTTCTAAG-3’ (WT amplicon 480 bp and floxed allele amplicon 350 bp); *Pten* floxed allele were forward 5’-AAAAGTTCCCCCTGCTGATTTGT-3’ and reverse 5’-TGTTTTTGACCAATTAAAGTAGGCTGT-3’ (WT amplicon 310 bp and floxed allele amplicon 435 bp). PCR conditions for ReddyMix (Thermo Scientific) were used according to manufacturer instructions with an initial denaturing step of 95°C for 2 minutes; 30 or 35 cycles of denaturing at 95°C for 25 seconds, annealing at 55°C for 35 seconds and extension at 72°C for 65 seconds, followed by a final extension at 72°C for 5 minutes. PCR products were separated on a 2% agarose gel and genotype determined by the absence or presence of expected amplicons.

*Red Blood Cell (RBC) Folate Measurement*

RBC folate was measured using a competitive immunoassay (Advia Centaur Folate assay) according to manufacturer instructions (Siemens Medical Solutions Diagnostics, Tarrytown, NY) and drawn from peripheral blood drawn via the retro-orbital sinus. Peripheral blood cell hematocrit level, needed to calculate RBC folate concentration, was determined using Hemavet 850 according to manufacturer instructions (CDC Technologies Inc., Oxford, CT).

Supplementary Table 1. (a) Number of dorsal root ganglia (DRG), (b) brachial plexi enlargement, and (c) sciatic nerve enlargement according to dietary folate intake

(a)
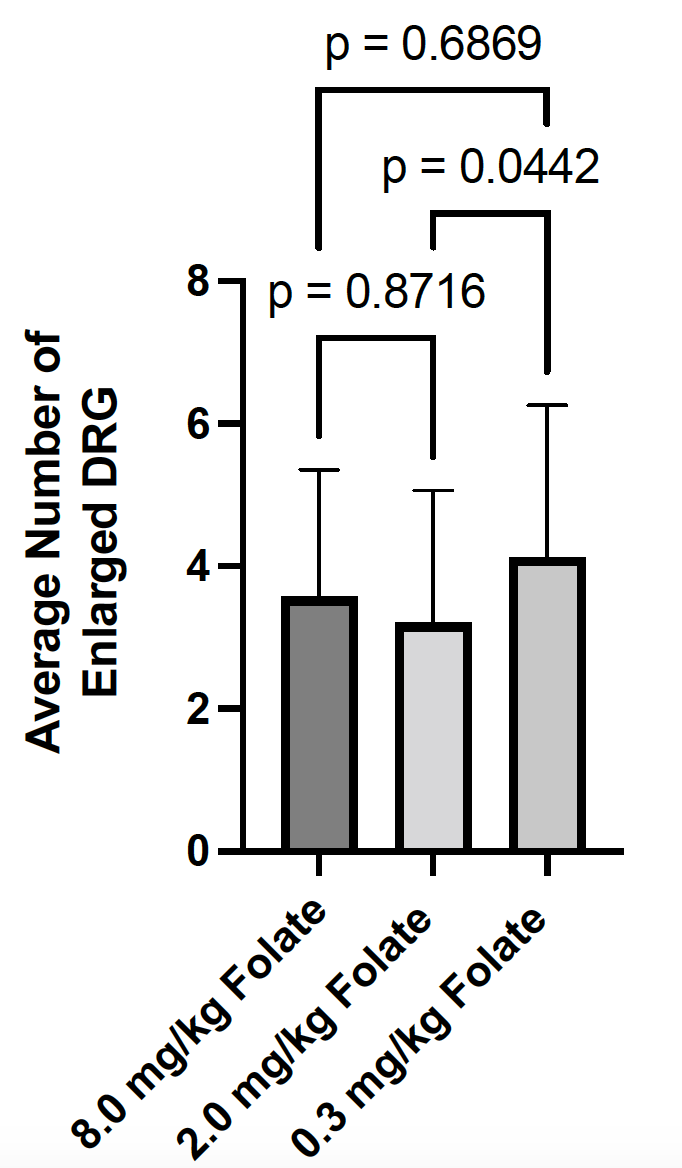
 (b)
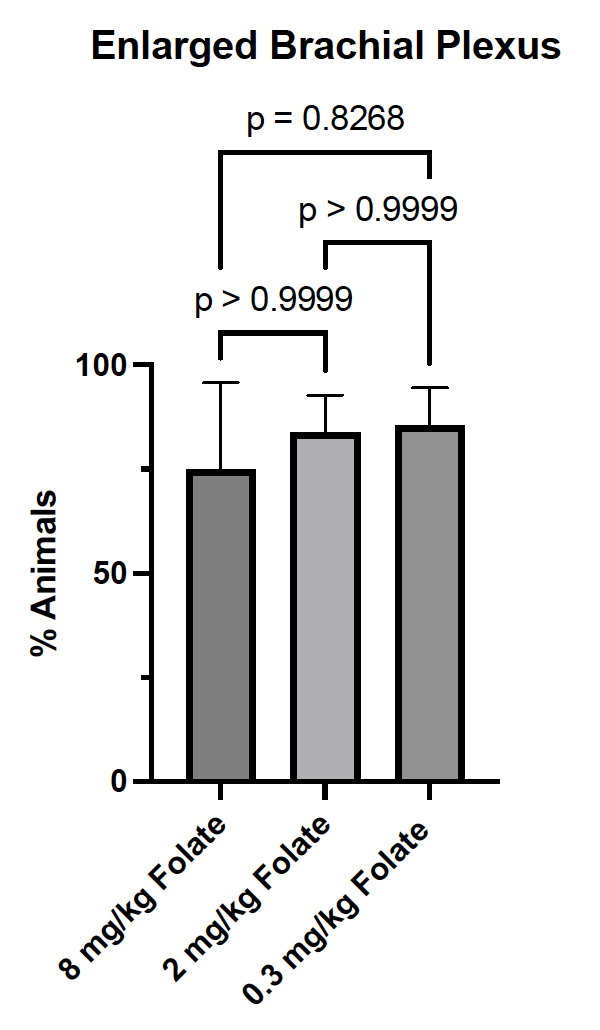
 (c)
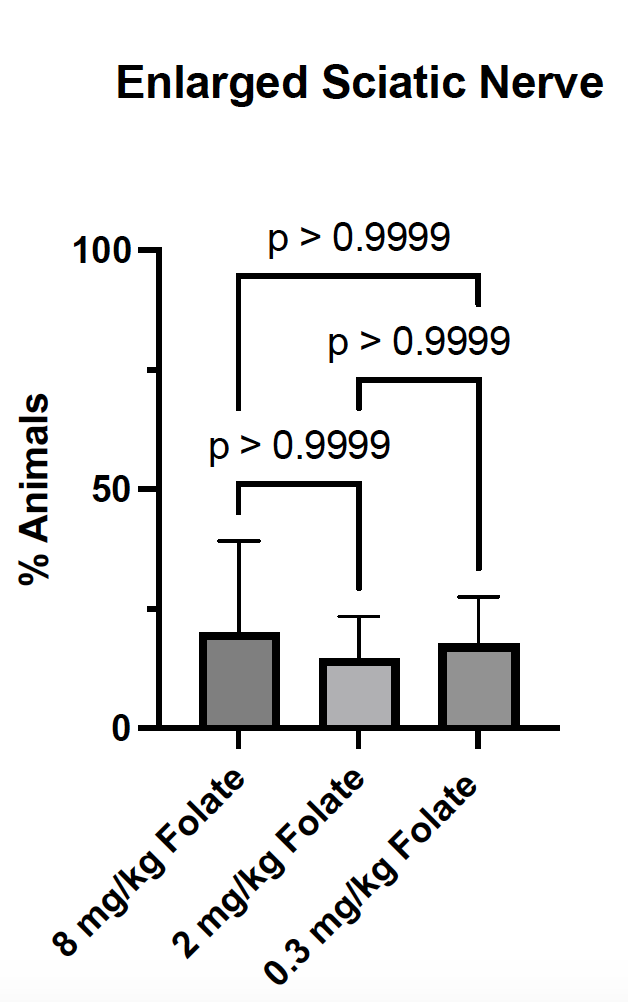
